# Effects of Altered Excitation-Inhibition Balance on Decision Making in a Cortical Circuit Model

**DOI:** 10.1101/100347

**Authors:** Norman H. Lam, Thiago Borduqui, Jaime Hallak, Antonio C. Roque, Alan Anticevic, John H. Krystal, Xiao-Jing Wang, John D. Murray

## Abstract

**Background:** Disruption of the synaptic balance between excitation and inhibition (E/I balance) in cortical circuits is a leading hypothesis for pathophysiologies of neuropsychiatric disorders, such as schizophrenia. However, it is poorly understood how synaptic E/I disruptions propagate upward to induce cognitive deficits, including impaired decision making (DM).

**Methods:** We investigated how E/I perturbations may impair temporal integration of evidence during perceptual DM in a biophysically-based model of association cortical microcircuits. Using multiple psychophysical task paradigms, we characterized effects of NMDA receptor hypofunction at two key synaptic sites: inhibitory interneurons (elevating E/I ratio, via disinhibition), versus excitatory pyramidal neurons (lowering E/I ratio).

**Results:** Disruption of E/I balance in either direction can similarly impair DM as assessed by psychometric performance, following inverted-U dependence. Nonetheless, these regimes make dissociable predictions for task paradigms that characterize the time course of evidence accumulation. Under elevated E/I ratio, DM is impulsive: evidence early in time is weighted much more than late evidence. In contrast, under lowered E/I ratio, DM is indecisive: evidence integration and winner-take-all competition between options are weakened. These effects are well captured by an extended drift-diffusion model with self-coupling.

**Conclusions:** Our findings characterize critical roles of cortical E/I balance in cognitive functions, the utility of timing-sensitive psychophysical paradigms, and relationships between circuit and psychological models. The model makes specific predictions for behavior and neural activity that are testable in humans or animals under causal manipulations of E/I balance and in disease states.

## Introduction

A central challenge in clinical neuroscience is to bridge explanatory gaps between the level of neurons and synapses, where pathophysiological mechanisms occur, and the level of cognitive processes, where behavioral symptoms are manifested and diagnosed. A burgeoning multidisciplinary approach to this challenge, computational psychiatry, leverages advances in theoretical neuroscience to investigate the impact of mechanistic disturbances on emergent brain function (1–3). Biophysically-based models of cortical circuits that implement cognitive functions can make dissociable predictions at the levels of behavior and neural activity arising from distinct synaptic-level perturbations (1,4).

Disruption in the synaptic balance between excitation and inhibition (E/I balance) in cortex is a leading hypothesis for pathophysiologies of neuropsychiatric disorders including schizophrenia (SCZ) (5–10) and autism spectrum disorder (ASD) (11–14). SCZ and ASD are associated with prominent deficits in cognitive function (15–18). It remains poorly understood how disruptions of E/I balance at the synaptic level propagate upward to the behavioral level, leading to specific deficits in cognitive computations.

A core component of cognitive function is decision making (DM), the deliberative process of forming a categorical choice. In many DM task paradigms, decisions are based on the accumulation of perceptual evidence over time. In one highly influential task paradigm, the subject must decide the net direction of random-dot motion (RDM) stimuli, which encourages DM based on the temporal integration of momentary perceptual evidence (19–23). Applied to clinical populations, RDM paradigms reveal impaired perceptual discrimination in SCZ (24–26) and ASD (27,28). In addition to potential dysfunction in upstream sensory representations (29), DM impairments may have contributions from dysfunction in evidence accumulation within association cortical circuits. In parietal and prefrontal association cortex, ramping neuronal activity reflects accumulated perceptual evidence, and activity crossing a threshold corresponds to decision commitment. These neurophysiological signatures have been interpreted in terms of the drift-diffusion model (DDM) from mathematical psychology (21).

At the level of neural mechanisms, studies in computational neuroscience have found that biophysically-based models of association cortical circuits can capture key behavioral and neurophysiological features during perceptual DM (30–33). DM function in the circuit model depends on strong synaptic interactions among excitatory pyramidal neurons and inhibitory interneurons. Recurrent excitation among pyramidal neurons, mediated by slow NMDA receptors (NMDARs), enables evidence accumulation. Feedback inhibition mediated by interneurons enables competition among neuronal populations and categorical decisions.

Here we characterized how disruptions in cortical E/I balance affect DM function in a previously validated spiking circuit model (30). We found that both elevated and lowered E/I ratio can impair DM function similarly, when assessed by standard psychometric performance. However, these E/I regimes make dissociable behavioral predictions under psychophysical paradigms that characterize the time course of evidence accumulation. These regimes could be well described by a drift-diffusion model (DDM) (21), which is modified so that integration is imperfect. This study therefore links synaptic disruptions to cognitive dysfunction in a well-established perceptual DM paradigm, and makes empirically testable predictions for DM behavior under elevated vs. lowered E/I ratio in association cortex.

## Methods and Materials

### Cortical Circuit Model

The biophysically-based circuit model, consisting of interconnected excitatory pyramidal neurons and inhibitory interneurons, represents a local microcircuit in association cortex (30). The circuit contains two populations of pyramidal neurons, each selective to evidence for a choice *A* or *B* (e.g., left vs. right). During stimulus presentation, the inputs to the populations represent momentary sensory evidence, reflecting the RDM stimulus coherence. 100%-coherence stimuli, corresponding to all dots moving coherently in one direction, are simulated by maximal (minimal) input to the preferred (anti-preferred) population. 0%-coherence stimuli, corresponding to dots moving incoherently with no net global motion, are simulated by equal input to both populations (**Figure 1A**). Stimulus input drives categorical, winner-take-all choice in the circuit. Due to inherent stochasticity from Poisson irregular spike trains received by all neurons, the model generates probabilistic choices whose proportions vary in a graded manner with coherence.

**Figure 1:**
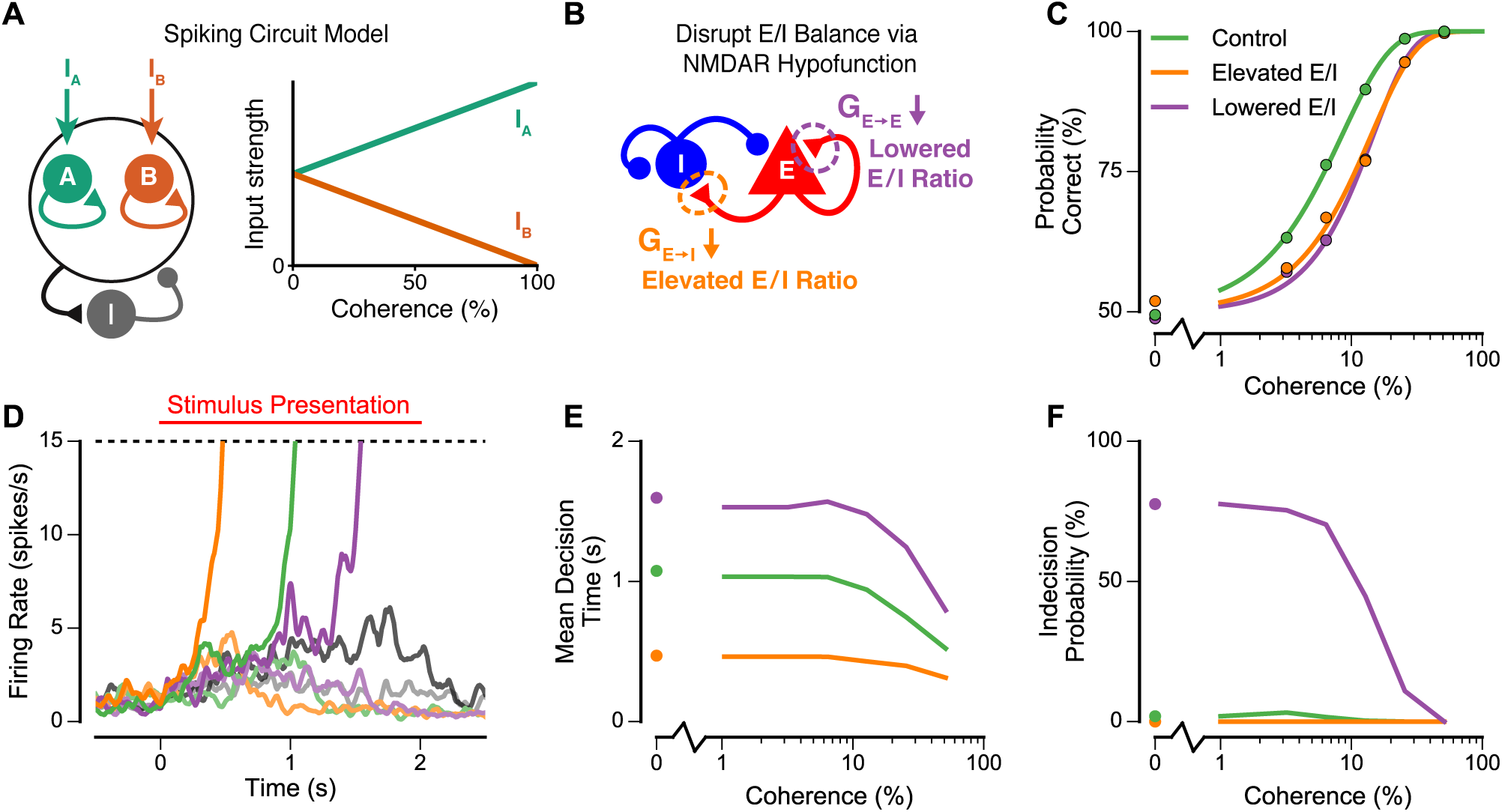
Model architecture and decision making (DM) performance under perturbations of excitation inhibition (E/I) balance. **(A)** Schematic circuit architecture. The model consists of recurrently connected excitatory pyramidal neurons (E) and inhibitory interneurons (I). The circuit contains two populations of pyramidal neurons which are each selective to one of the two stimuli (A and B). Within each pyramidal-neuron population there is strong recurrent excitation. The two populations compete via feedback inhibition mediated by interneurons. **(B)** E/I perturbations via NMDAR hypofunction on two synaptic sites. NMDAR hypofunction on inhibitory interneurons (reduced *G*_*E*→*I*_) weakens the recruitment of feedback inhibition, which elevates the E/I ratio via disinhibition. NMDAR hypofunction on excitatory pyramidal neurons (reduced *G*_*E*→*E*_) weakens recurrent excitation, which lowers the E/I ratio. **(C)** DM performance as quantified by the psychometric function, i.e., the proportion of correct choices as a function of coherence. Both perturbations, elevated and lowered E/I ratio, degrade performance relative to the control circuit. **(D)** Neuronal activity during DM. During stimulus presentation, the two neural populations (shown here in dark and light shading) compete. When one population crosses the firing-rate threshold (15 spikes/s), the corresponding behavioral choice is selected. The elevated-E/I circuit ramps more quickly than the control circuit, and the lowered-E/I circuit ramps more slowly. These example curves show the trial with median reaction time. The curves in gray show an example of an indecision trial from the lowered-E/I circuit, in which neither population crosses the firing-rate threshold. **(E)** Mean decision times, as a function of stimulus coherence. The decision time is the time from stimulus onset until one population first crosses the firing-rate threshold, on a trial in which this threshold is crossed. The elevated-E/I circuit and lowered-E/I circuit have shorter and longer decision times, respectively, compared to the control circuit. **(F)** The proportion of indecision trials, as a function of stimulus coherence. An indecision trial is defined as a trial in which neither neural population crosses the firing-rate threshold during the stimulus presentation. In two-alternative forced choice (2AFC) task paradigms, the choice is then selected randomly. The lowered-E/I circuit shows a high proportion of indecision trials for low coherence, which drives the performance deficit in this regime.

All tasks simulated follow a two-alternative forced choice (2AFC) paradigm. A categorical decision in the circuit is made when the corresponding population crosses a threshold firing rate, as observed in neuronal activity (20). If neither population crosses the threshold within 2 s following stimulus offset, then choice is assigned randomly, as implemented previously for 2AFC (30).

We perturbed E/I balance in the model bidirectionally through hypofunction of ND-MARs at two sites: on inhibitory interneurons (I-cells), or on excitatory pyramidal neurons (E-cells) (**Figure 1B**). NMDAR hypofunction on I-cells (reduced *G_E→I_*) results in elevated E/I ratio via disinhibition, whereas NMDAR hypofunction on E-cells (reduced *G_E→E_)* results in lowered E/I ratio. We selected NMDAR hypofunction as a mechanism to alter E/I ratio because it is a leading hypothesis in the pathophysiology of SCZ (6–8), and NMDAR antagonists such as ketamine provide a leading pharmacological model of SCZ (5,34). In doing so, the presented simulations provide a directly empirically testable set of predictions for future pharmacological and clinical studies (35). For default perturbations, in the elevated-E/I circuit, *G_E→I_* is reduced by 3%. Conversely, in the lowered-E/I circuit, *G_E→E_* is reduced by 2%. These magnitudes preserve stability of the low-activity baseline state and high-activity memory state.

### Psychophysical Tasks

We used four psychophysical 2AFC task paradigms to characterize DM function, one standard and three which characterize the time course of evidence accumulation, which are all adapted from primate electrophysiological studies (for details, see **Supplement 1**):

1. In the “standard” task paradigm, stimulus is presented for a fixed 2-s duration at a constant coherence level, and the coherence varies trial-by-trial (19,30). The psychometric function, giving the percent correct as a function of coherence, defines the discrimination threshold as the coherence eliciting 81.6% correct (36).
2. In the “psychophysical kernel” task paradigm, the 2-s stimulus duration is subdivided into multiple time bins (0.05-s each), whose coherence values are sampled independently from a zero-mean, uniform distribution of coherences (Figure 3A) (37). The psychophysical kernel is computed as a choice-triggered average to quantify the contribution of each time bin to the resulting behavioral choice (36–39).
3. In the “pulse” task paradigm, the stimulus is the same as that of the standard task, except that a brief (0.1-s) pulse of additional coherence (±15%) is applied at a variable onset time (32,36,40). The pulse induces a shift of the psychometric function according to pulse coherence, and we characterize dependence of this shift on pulse onset time.
4. In the “duration” task paradigm, the stimulus is the same as that of the standard task, except that the stimulus duration is varied trial-by-trial (30,36). For a given duration, we measure the psychometric function and obtain a discrimination threshold. We then and characterize dependence of the threshold on stimulus duration.

**Figure 3:**
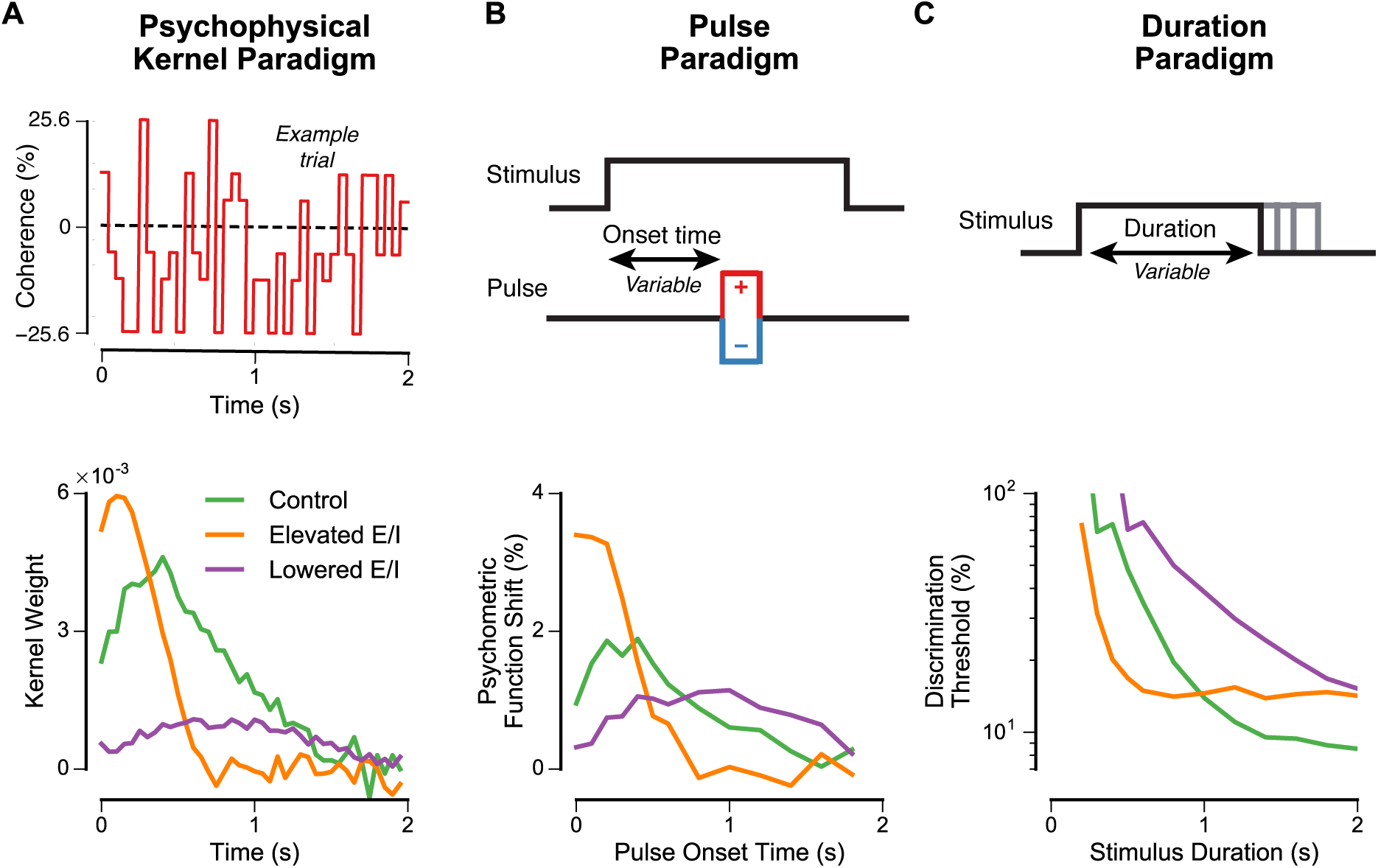
DM task paradigms that characterize the time course of evidence accumulation can dissociate elevated vs. lowered E/I ratio. **(A)** Top: Psychophysical kernel (PK) paradigm. This paradigm uses randomly time-varying stimuli to characterize how much weight a given time point has on the choice. Bottom: PK weights as a function of time during the stimulus, for the three E/I regimes (see also **Figure S2A–C**). Stimuli at time points with a large PK weight are those with a large influence on the choice, whereas stimuli at time points with PK weight near zero have little effect on choice. In the control circuit, the PK weight shows an initial rise followed by a gradual decrease over time. Relative to the control circuit, the elevated-E/I circuit heavily weights early time points but assigns little weight to late time points. Relative to the control circuit, the lowered-E/I circuit has a flattened and weakened profile for the PK weight, indicating less impact of all stimuli on choice. **(B)** Top: Pulse paradigm. This paradigm uses a brief pulse of additional perceptual evidence at different onset times to characterize the degree to which it shifts the psychometric function. Bottom: Shift in the psychometric function as a function of pulse onset time, for the three E/I regimes (see also **Figure S2D–F**). Similar to the PK, the control circuit shows an initial rise and then a gradual decrease in the magnitude of the shift as a function of the pulse onset time. Relative to control, in the elevated-E/I circuit the pulse has a stronger impact at early onset times, but less impact at later onset times. The lowered-E/I circuit shows a flattened profile of the shift, with greater impact at late onset times. **(C)** Top: Duration paradigm. This paradigm varies the duration of the stimulus presentation trial-by-trial. For a given stimulus duration, the discrimination threshold is measured from the psychometric function (see also **Figure S2G–I**). Bottom: Discrimination threshold as a function of stimulus duration, for the three E/I regimes. Discrimination thresholds decrease with longer stimulus durations. The key property is the duration at which the threshold plateaus, indicating limits to temporal integration performance. For the elevated-E/I circuit, the threshold plateaus at shorter durations relative to control. For the lowered-E/I circuit, the threshold does not show a strong plateau up to the maximum stimulus duration of 2 s.

### Extended Drift-Diffusion Model with Self-Coupling

We tested whether behavioral effects of disrupted E/I balance in the circuit model can be captured by an extended DDM (41). We hypothesized that E/I circuit alterations would correspond to alterations of the DDM integration process. Therefore the extended DDM includes a self-coupling term *λ*, so that instead of perfect integration (*λ* =0), the integration process can be leaky (*λ*<0) or unstable (*λ*>0). We simulated the extended DDM via a Fokker-Planck formalism (see **Supplement 1** for details).

## Results

### E/I Balance Affects Decision Making in a Cortical Circuit Model

We first characterized DM performance via the psychometric function in the standard task paradigm, in which the stimulus has a constant coherence that varies across trials and a fixed duration. The probability of correct choice increases monotonically with higher coherence (**Figure 1C**). We found that both the elevated and lowered E/I ratio conditions yielded worse performance (higher discrimination threshold) relative to control. Therefore, performance in the standard task alone is insufficient to dissociate these neurophysiological regimes.

To gain insight into modes of dysfunction under these two manipulations, we next examined dynamics of neural activity during DM. **Figure 1D** shows representative single-trial activity traces for zero-coherence stimuli. In the elevated-E/I circuit, DM-related ramping of activity to threshold is much faster relative to control (**Figure 1D,E**). This suggests that DM performance is impaired because the decision process is based on less of the available perceptual evidence and the signal-to-noise ratio is not improved by longer temporal integration. In contrast, in the lowered-E/I circuit, neuronal activity fails to reach threshold on a greater fraction of trials, resulting in more random choices on low-coherence trials (**Figure 1D,F**). Therefore, DM in the elevated- and lowered-E/I circuits can be characterized as ‘impulsive’ and ‘indecisive,’ respectively. These findings show that the similarly impaired psychometric performance is due to distinct mechanisms in the two regimes.

To explicitly examine the circuit’s dependence on E/I ratio, we parametrically decreased both the NMDAR conductances on interneurons (*G_E→I_*) and on pyramidal neurons (*G_E→E_*), while characterizing DM performance via the psychometric function (**Figures 2A**,**S1**). For relatively small perturbations tested here, if *G_E→I_* and *G_E→E_* are reduced together in a certain proportion, DM performance is unaltered because E/I balance is maintained. Large parallel reductions can impair DM function, because winner-take-all competition depends on strong recurrent excitation and inhibition (31). Reduction of *G_E→l_* in greater proportion elevates E/I ratio and degrades performance due to impulsive DM. In contrast, reduction of *G_E→E_* in greater proportion lowers E/I ratio and degrades performance due to decisive DM. DM performance thus exhibits an “inverted-U” dependence on E/I ratio (**Figure 2B**). These findings indicate that E/I ratio is a crucial effective parameter for DM function in the circuit, rather than the absolute strength of one synaptic connection alone.

**Figure 2:**
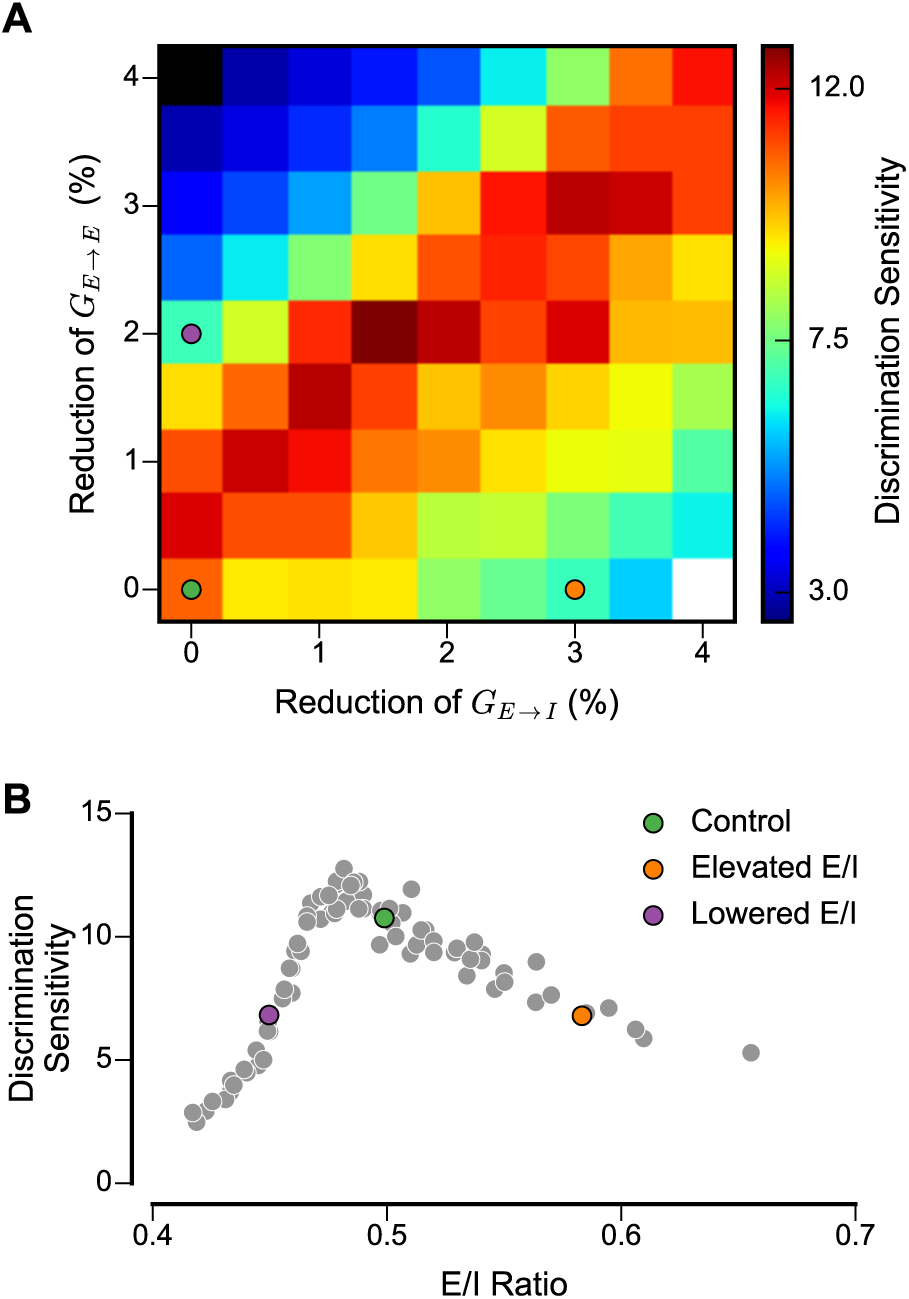
The parameter space of NMDAR hypofunction highlights the importance of E/I balance for DM function. **(A)** Sensitivity, defined as the inverse of the threshold of the psychometric function for the standard task, varies as a function of the NMDAR conductance on inhibitory interneurons (G_*E*→*I*_) and on excitatory pyramidal neurons (*G_*E*→*E*_*). Sensitivity can be maintained with a proportional decrease in both *G_*E*→*I*_* and *G*_*E*→*E*_. In the white region at the bottom right corner, the low-activity baseline state is destabilized, due to strong disinhibition. In the black region at the top left corner, the high-activity memory state is destabilized, due to insufficient recurrent excitation. **(B)** Inverted-U dependence of DM performance on E/I ratio. E/I ratio is defined as the ratio of the net recurrent excitatory current (AMPA and NMDA) to net inhibitory current (GABA_a_), for pyramidal neurons. Gray circles represent the combinations of {*G_E→I_*,*G_E→E_*} parameter values shown in **(A)**.

The above findings suggest that standard psychometric functions, as measured in clinical populations (24–28), are not well suited to dissociate among distinct forms of DM impairment potentially related to different underlying pathophysiological states. Are there behavioral tasks that can dissociate between these distinct DM impairments induced by elevated vs. lowered E/I ratio? Based on DM-related neural activity (**Figure 1D-F**), we hypothesized that these regimes would make dissociable predictions for the time course of evidence accumulation. Next, we describe three task paradigms, grounded in electrophysiological studies of perceptual DM, that characterize this time course (**Figures 3**,**S2**).

### Psychophysical Kernel Paradigm

One thorough method to characterize the time course of evidence accumulation is through the psychophysical kernel (PK) task paradigm, which uses randomly time-varying stimulus to quantify the weight a given time point has on behavioral choice (**Figures 3**,**S2A-C**). Stimuli at time points with a large PK weight have a large influence on choice, whereas stimuli at time points with PK weight near zero have little impact on choice. We found that for the control circuit, the PK exhibits an initial rise and then decay, which is qualitatively consistent with PKs measured in a number of primate electrophysiology experiments during perceptual DM (36–38). The decline of the PK with time reflects bounded accumulation (36): after the circuit reaches a decision state, stimuli at subsequent time points do not affect the choice on that trial. The PK is therefore shaped by the distribution of threshold crossing times.

Relative to control, the PK for the elevated-E/I circuit assigns much more weight to early time points and much less weight to late time points. For the lowered-E/I circuit, the PK is generally flattened and has lower weights. In this regime, the choice is generally less driven by evidence (i.e., the integral of the PK is low). These PK findings are consistent with characterizations of impulsive vs. indecisive DM for elevated vs. lowered E/I ratio, respectively. In contrast to the standard task paradigm, these two E/I perturbations make dissociable behavioral predictions in the PK task paradigm.

### Pulse Paradigm

Another psychophysical task paradigm to directly test for differential effects of early vs. late evidence measures the impact of a brief pulse of additional perceptual evidence as a function of its onset time (32,36,40). The impact of the pulse can be quantified by a horizontal shift in psychometric function, according to pulse coherence. Kiani et al. found that this pulse-induced shift decreased at later pulse onset times (36).

We found that the time dependence of this pulse-induced shift mirrors the time course of the PK (**Figures 3B**,**S2D-F**). In the control circuit, the magnitude of the shift increased over very early onset times and then decreased over later onset times. Relative to control, the elevated-E/I circuit showed a stronger shift for early onset times, but the psychometric function was much less sensitive to pulses at later onset times. In contrast, the lowered-E/I circuit showed a flattened pattern.

### Duration Paradigm

The PK and pulse paradigms directly reveal the time course of evidence accumulation, and its bounded nature, in the circuit model. If integration were perfect and unbounded, then increasing the stimulus duration in the standard task paradigm should always reduce the discrimination threshold. However, bounded integration implies that this improvement should plateau at long durations, as observed empirically (36). The key feature in this characterization is the duration at which the threshold plateaus without further improvement, reflecting the timescale of evidence accumulation.

Varying the stimulus duration trial-by-trial, we found that the duration of plateau differs substantially across the E/I regimes, in a manner consistent with the PK and pulse findings (**Figures 3C**,S2G-I). In the elevated-E/I circuit, performance plateaus at a shorter duration, relative to control. Interestingly, the elevated-E/I circuit performs better than control for short stimulus presentations, because for short durations, the control circuit fails to make an internal choice on a greater fraction of trials (see Discussion). In contrast, the lowered-E/I circuit shows high discrimination thresholds which decline without plateauing up to 2-s durations.

### Comparison to an Upstream Sensory Coding Deficit

The above analyses consider dysfunction within a DM circuit. A third potential cause of DM impairment is dysfunction in upstream sensory areas which transmit the perceptual evidence signals to downstream DM circuits. To test the impact of this mechanism, we characterized the control circuit under sensory inputs that showed a weakened modulation by stimulus coherence (**Figure S3**). We found that an upstream sensory coding deficit can similarly impair discrimination in the standard task paradigm. However, it makes dissociable predictions for behavioral measures in the other task paradigms. Although this deficit can scale the amplitudes of the measures, their overall dependences on stimulus timing, reflecting evidence accumulation, are preserved.

### Comparison to an Extended Drift-Diffusion Model

We next sought to relate the effects of E/I imbalance in the circuit model to a highly influential theoretical framework for 2AFC DM, the drift-diffusion model (DDM) (42). The DDM describes the dynamics of a decision variable that accumulates noisy evidence over time. An internal decision is made when the decision variable crosses a corresponding bound. In the standard DDM, the temporal integration process is perfect, i.e., its memory timescale is infinite and the decision variable is simply the time integral of its evidence input.

We hypothesized that E/I alterations may map onto changes in the temporal integration process itself. To capture such effects, we extended the DDM to include a self-coupling term *λ* (**Figure 4A**) (see **Supplement 1**) (41,43). *λ*=0 corresponds to the perfect integrator of the standard DDM, with an infinite time constant for memory (*τ =* |*λ*|^−1^). *λ*<0 corresponds to a leaky integrator, with a finite time constant for memory. *λ*>0 corresponds to an unstable integrator, which has an intrinsic tendency to diverge away from zero (**Figures 4B**,**S4**) (43). We predicted that DM effects of elevated and lowered E/I ratio could be well captured by unstable and leaky integration, respectively.

**Figure 4:**
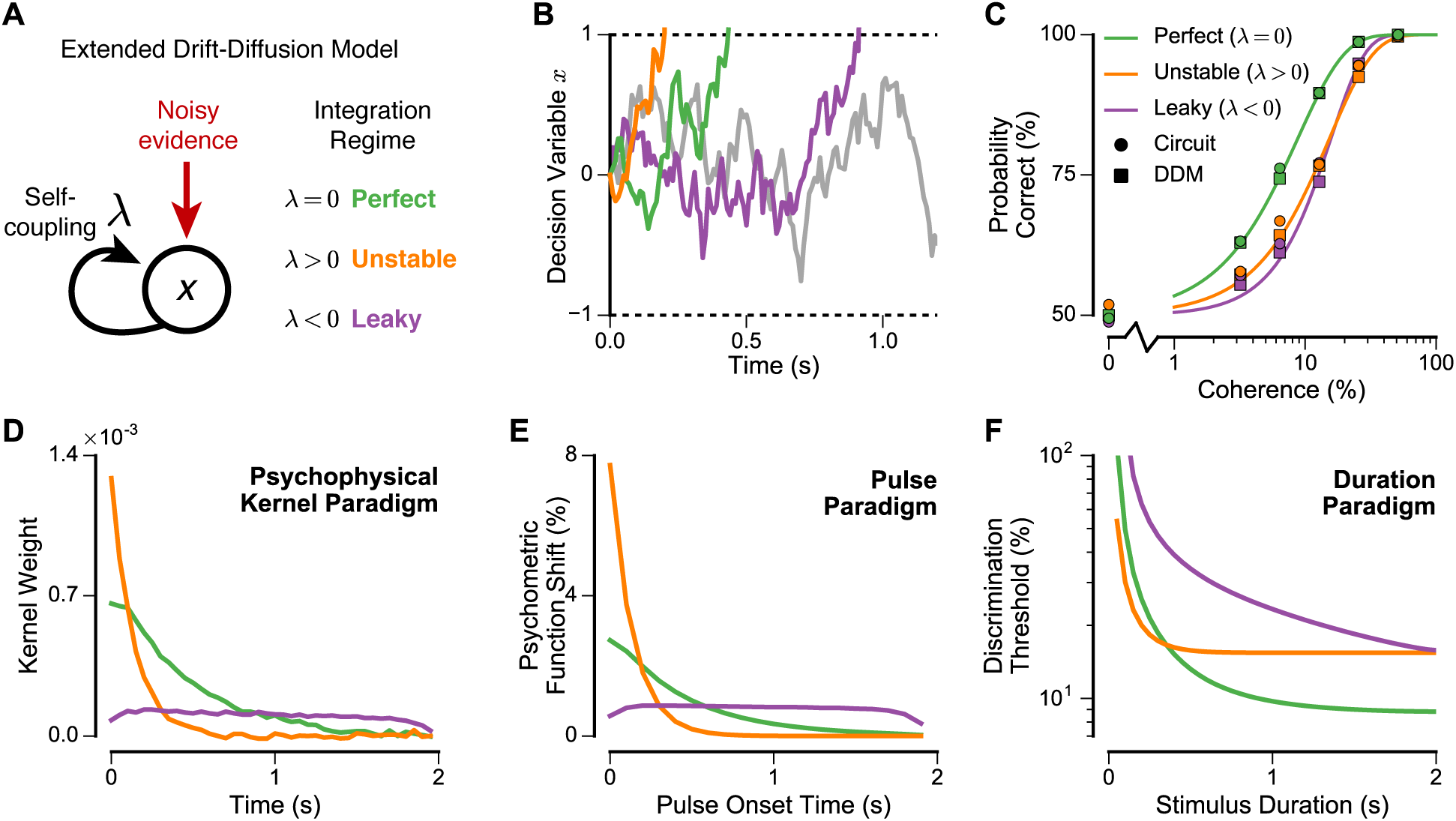
An extended drift-diffusion model (DDM) with a self-coupling term captures the effects on DM of elevated vs. lowered E/I ratio. (**A**) Schematic for the extended DDM. We extended the standard DDMto include a self-coupling term λ, such that the input to the decision variable *x* includes a term *λx*. λ = 0 corresponds to a perfect integrator, which is the standard DDM. *λ* < 0 corresponds to a leaky integrator with finite-timescale memory. *λ* > 0 corresponds to an unstable integrator with a tendency to diverge from 0. (**B**) Example trajectories of the decision variable *x* for the three self-coupling regimes. In the DDM, the decision variable performs stochastic accumulation of noisy evidence, moving it toward option *A* (positive) or option *B* (negative). When it crosses the corresponding threshold, the choice is selected. In the unstable regime the decision variable rapidly diverges from 0 and crosses a threshold, whereas in the leaky regime the threshold is crossed much later. These example traces show the trials with median reaction time. The trajectory in gray shows an example of an indecision trial from the leaky regime. (**C**) Psychometric functions for the three self-coupling regimes. The self-coupling parameters for the unstable and leaky regimes are quantitatively fitted to psychometric performance of the elevated-E/I and lowered-E/I circuits, respectively (see **Supplement 1 and Figure S6**). Fitting yielded *λ* = 7.0 s^−1^ (*τ* = 0.14 s, time constant of unstable growth) for the elevated-E/I circuit and *λ* = −7.7 s^−1^ (*τ* = 0.13 s, time constant of leak) for the lowered-E/I circuit. Circles and squares mark data from the circuit model and extended DDM, respectively, and the curves mark the psychometric function fitted to the extended DDM. (**D–F**) Psychometric characterization for self-coupling regimes, using fitted parameter values, for the psychophysical kernel paradigm (**D**), pulse paradigm (**E**), and duration paradigm (**F**) (see also **Figures S6, S7**).

To test these hypotheses, we first fitted a perfect integrator (*λ*= 0) model to the control circuit, and then fitted the *λ* parameter to the two disrupted conditions, using behavioral and neural features in the standard task paradigm (**Figures 4C**,**S5**) (see **Supplement 1**). In line with our predictions, fitting *λ* yielded a large positive value for the elevated-E/I circuit, corresponding to a highly unstable integrator, and a large negative value for the lowered-E/I circuit, corresponding to a highly leaky integrator. We then tested these extended DDMs on the three paradigms that probe the time course of evidence accumulation. Remarkably, we found that these two regimes of self-coupling could well capture key aspects of DM behavior across all paradigms (**Figures 4D-F**,**S6**,**S7**). A notable difference occurs at early time points, where the extended DDM is sensitive to stimuli immediately following stimulus onset (see Discussion).

## Discussion

In this study, we characterized the role of E/I balance in DM in a biophysically-based cortical circuit model. Elevated or lowered E/I ratio can degrade DM performance, following an inverted-U dependence, but for different underlying reasons: impulsive and indecisive DM for elevated and lowered E/I ratio, respectively. We found that task paradigms can dissociate these regimes by characterizing the time course of evidence accumulation.

Our circuit model is grounded in extensive studies in monkeys performing perceptual DM with RDM stimuli, which have characterized psychophysical behavior and neuronal activity from sensory and association cortex (19,20,36,40,44). For RDM stimuli, early visual area MT is critical for sensory processing of motion signals. Electrophysiological recordings during RDM DM have found that MT represents the momentary motion evidence (21,44). Downstream parietal and prefrontal association cortical areas represent the accumulation of evidence in the form of a decision variable (19–21).

### Relation to Clinical Findings

Applied to neuropsychiatric populations, motion discrimination tasks with RDM stimuli have revealed impaired perceptual discrimination (i.e., higher discrimination thresholds) in SCZ (24–26) and ASD (27,28), both of which are associated with disrupted cortical E/I balance (5–14). Prior psychophysical studies in SCZ and ASD primarily interpreted impaired discrimination as reflecting dysfunction in upstream early visual cortex (29), such as area MT. Present findings suggest a potential complementary source of DM impairment from downstream association cortex involved in evidence accumulation. The proposed task paradigms dissociate different forms of DM impairment from an upstream sensory deficit, and can reveal concurrent alterations in sensory and association circuits.

Our use of the terms “impulsive” and “indecisive” refers specifically to the time course of evidence accumulation in DM, and thus our model may not relate to clinically relevant forms of behavioral impulsivity (45), nor does the indecisive regime imply failure of behavioral response in 2AFC tasks. It remains unknown how processes of evidence accumulation during perceptual DM, described here, may relate to other clinical abnormalities in DM. For instance, in a sequential perceptual inference task, SCZ subjects exhibit “jumping to conclusions,” basing decisions on much less perceptual evidence relative to controls (46).

### Testing Model Predictions

The circuit model makes dissociable predictions for DM behavior arising from distinct synaptic perturbations to E/I balance, which can be tested in animal models and in humans. Animal models allow for causal perturbation of E/I balance in association cortex during DM. Optogenetic, pharmacological, or developmental manipulations can elevate or lower cortical E/I ratio in a graded manner and affect cognitive behavior (13,39,47). For instance, in a rodent perceptual DM study, agonism of GABA_A_ receptors in a parietal DM-related area decreased the PK amplitude (39).

The proposed psychophysical task paradigms, which characterize the time course of evidence accumulation, have the potential to elucidate distinct modes of DM impairment in neuropsychiatric disorders such as SCZ and ASD (24–28). The higher discrimination threshold, studied in the standard task, may have different underlying DM deficits across clinical populations. The neurophysiological basis of the circuit model allows interpretation of dissociable behavioral regimes in terms of underlying differences in cortical E/I balance. There is growing evidence that individual differences in E/I ratio of association cortex contribute to differences in cognitive function (48–50). Quantitative behavioral analysis, informed by model hypotheses, can allow strong inference of circuit regimes (51).

In healthy humans, cortical E/I balance can be perturbed pharmacologically, e.g., via subanesthetic administration of ketamine (4,35). Perceptual DM with RDM stimuli has been studied during pharmacological manipulation in humans. For instance, one study found that psilocybin impairs perceptual discrimination with RDM stimuli (52). The effects of neuromodulators, such as dopamine and serotonin, can be incorporated into these biophysically-based circuit models and related to cognitive functions (53,54). Multiple mechanisms that alter E/I ratio may converge in their neural and behavioral consequences (4).

### Comparison to Drift-Diffusion Models

An important goal supporting computational psychiatry is to connect models that operate at different levels of abstraction (1–3), such as the synaptic-level circuit models and the psychological-level DDMs considered here. Applications of the DDM typically assume perfect integration, and capture deviations from standard DDM behavior by introducing collapsing bounds or urgency signals (55). Our findings suggest the utility of incorporating a self-coupling term into an extended DDM to capture the behavioral impact of leaky or unstable integration (43,56,57). Depending on task demands, it may be functionally beneficial to flexibly modulate the integration timescale, e.g., for dynamic stimuli whose reliabilities vary with time (58,59). Future studies could probe how flexible adaptation of integration processes may be impaired in clinical populations.

The extended DDM was able to capture key features of the elevated-E/I and lowered-E/I circuits during psychophysical paradigms characterizing the time course of evidence accumulation. Biophysically-based circuit models are highly computationally intensive to simulate, which limits their practical application to fit empirical data. In contrast, the extended DDM is computationally tractable when simulated using the Fokker-Planck formalism. The extended DDM thereby enables tractable fitting to empirical and modeling-generated psychometric choice data. Therefore, a potentially fruitful strategy using these psychophysical tasks may be to fit an extended DDM to empirical choice (60). The circuit model, establishing links between imperfect integration in the extended DDM and alterations in cortical E/I ratio, provides mechanistic insight into the dependence of cognitive impairments on underlying neurophysiology.

The task paradigms reveal a notable difference between the circuit model and the extended DDM. In the circuit, sensitivity to the stimulus is relatively low at very early time points and rises to maximum sensitivity. This is because following stimulus onset, neural activity takes time to approach a coherence-sensitive integrative state (31). In contrast, the extended DDM begins integration immediately following stimulus onset. However, this time to reach an integrative state is not a fixed property, as external inputs can place the circuit closer to the integration state (31,32).

### Implications for Task Design

In this study we considered ‘fixed duration’ task paradigms, in which the experimenter controls the duration of stimulus presentation. Perceptual DM can also be studied in ‘reaction time’ paradigms, in which the subject is free to respond at any time during stimulus presentation (20). The reaction time is thought to vary with the internal decision time, but also with a ‘non-decision’ time encompassing other stages of sensorimotor processing. Interpretation of reaction times in terms of underlying decision times may be complicated for clinical or pharmacological comparisons. Even in simple choice tasks, patients with SCZ exhibit reaction times that are slower and more variable (61,62). Similarly, subanesthetic administration of ketamine lengthens reaction times (63,64). To avoid these potential confounds, we focused on fixed duration paradigms that can characterize the time course of evidence accumulation, rather than infer it from reaction-time variation. Among the paradigms considered, we propose the pulse paradigm may be best suited for studies with clinical populations, because the PK paradigm typically requires many trials with near-chance performance (37).

### Future Directions

Biophysically-based circuit models, which can link synaptic perturbations to behavior, provide a powerful proof-of-principle platform to explore potential compensations mediated by pharmacological interventions. Following a perturbation at one site in the circuit, cortical E/I balance could be restored through compensations at other sites, through glutamatergic, GABAergic, or neuromodulatory modulations (4). The model suggests that E/I balance is the crucial parameter that determines DM performance. Therefore, compensations that restore E/I balance can potentially ameliorate observed DM deficits.

## Acknowledgements

Funding was provided by NIH grants DP5OD012109 (AA), R01MH062349 (X-JW), and TL1 TR000141 (JDM), and CNPq Science Without Borders grant #202853/2014 (TB). The authors declare no competing conflicts of interest.

## Supplemental Information

### Spiking Circuit Model

The spiking circuit model is based on the model of Wang (1), with only minor changes as reported below. All other details of the circuit model are described in Ref. (1).

The circuit is composed of *N_E_* = 1600 excitatory neurons and *N_I_ =* 400 inhibitory neurons, all simulated as leaky integrate-and-fire neurons. Recurrent excitatory connections are mediated by both NMDA and AMPA conductances. Recurrent inhibition is mediated by GABA_a_ conductances. Background and stimulus inputs are mediated by AMPA conductances with Poisson spike trains.

For decision making (DM) between two choices, we separate two non-overlapping groups of excitatory neurons, each of size *N_E,A_ = N_E,B_ =* 240, which correspond to two choices *A* and *B*. The two groups compete with each other via lateral inhibition mediated by interneurons. The remaining excitatory neurons are non-selective. The connectivity pattern between excitatory and inhibitory neurons, and among inhibitory neurons, are unstructured. The connectivity pattern among excitatory neurons follow the “Hebbian” form as in Ref. (1), where recurrent projections to neurons in the same group have a stronger synaptic strength (enhanced by a factor of *w_+_ >* 1). To preserve the total recurrent excitatory projections, the synaptic strength to neurons in the competing group as well as to non-selective neurons are modified by a factor of *w_−_ =* 1 − *f (w −* 1)/(1 − *f*), for *f = N_E1_*/*N_E_*= 0.15. The connections from non-selective neurons to all excitatory neurons are unstructured.

Stimulus-related signals from an upstream area produce inputs to the two neuron groups in the form of Poisson spikes, whose differential spike rates represent momentary sensory evidence in experiments. Inhibitory interneurons and non-selective excitatory neurons receive no sensory inputs. To simulate the representation of motion signals from a random-dot motion (RDM) stimulus in area MT, spike rates for sensory inputs to the two groups (*μ_A,B_*) are given by:

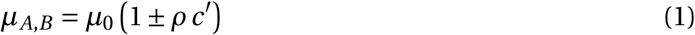

where *μ*_0_ is the overall stimulus strength, *ρ* is the upstream modulation parameter set to 1 by default, and *c′* is the stimulus coherence. A choice is selected when either excitatory neuron group reaches a threshold population firing rate of 15 Hz. If neither group crosses the threshold within 2 s following stimulus offset, then the choice is assigned randomly, as done previously for two-alternative forced choice (2AFC) task paradigms (1). The output spike data from the circuit model simulations are used to construct firing rates for each neuron group, via a casual exponential filter with a time constant of *τ*_*filt*_ = 20 ms. The firing rate is then used for further analyses such as threshold crossing.

We made the following minor adjustments to the original parameters of Ref. (1), to enhance the stability of the baseline and persistent-activity states when subject to E/I perturbations. E/I perturbations were implemented through reduction of the recurrent NMDA conductance either on inhibitory neurons (*G_E→I_*) or on excitatory neurons (*G_E→E_*). To stabilize the persistent-activity state against reductions of *G_E→E_*, we increased *w_+_* to 1.84 from 1.70. To stabilize the baseline state against reductions of *G_E→I_*, we reduced the external AMPA conductance (*g_ext_,AMPA*) to 2.07 nS from 2.1 nS. Finally, we changed the stimulus strength *μ*_0_ to 38 Hz from 40 Hz. For our default synaptic perturbation magnitudes, *G_E→I_* is reduced by 3% in the elevated-E/I circuit, and *G_E→E_* is reduced by 2% in the lowered-E/I circuit. These perturbation magnitudes preserve the stabilities of the low-activity baseline state and the high-activity memory state.

#### Stability Criteria

To determine the stability of the low-activity spontaneous state, for various perturbations to *G_E→E_* and *G_E→I_* (**Figures 2**,**S2**), we performed 10 sets of 5-s simulations for each conditions of *G_E→E_* and *G_E→I_* perturbation, with no evidence sensory inputs (**equation 1**) provided to any neurons. A circuit is deemed unstable if the spontaneous state reliably destabilizes in the majority of the simulations, such that the threshold population firing rate of 15 Hz is crossed by a group of excitatory neurons; otherwise the circuit is considered stable.

We also tested stability of the high-activity memory state, which enables the circuit to store the choice signal internally, through persistent activity during the delay, after a choice has been selected by threshold-crossing. For various perturbations to *G_E→E_* and *G_E→I_* (**Figure 2**,**S2**), we simulated 500 trials applying the 2-s, 0% coherence stimulus used in the standard task. For trials with threshold crossing, if the high-activity memory state is consistently maintained 2 s after stimulus presentation, the circuit is considered stable. Otherwise the circuit is considered unstable.

#### E/I Ratio Calculation

The E/I ratio in the spiking circuits with various perturbations to *G_E→E_* and *G_E→I_* (**Figure 2**, **S2**) are obtained from 10 sets of 5-s simulations for each conditions of *G_E→E_* and *G_E→I_* perturbation, in the stable baseline state with no sensory inputs. In the 5 s, the recurrent AMPA, NMDA, and GABA inputs to an excitatory neuron group is recorded. The E/I ratio is defined here as the total recurrent excitatory inputs divided by the recurrent inhibitory inputs.

### Psychophysical Task Paradigms

#### Standard Task Paradigm

In the standard task paradigm, a constant-coherence stimulus is applied for a fixed duration of 2 s. The coherence varies trial-by-trial from the set of {0%, 3.2%, 6.4%, 12.8%, 25.6%, 51.2%}. The psychometric function, giving the probability of a choice for option *A* as a function of coherence *c*′ (*P*(*c*′)), is fit by the functional form:

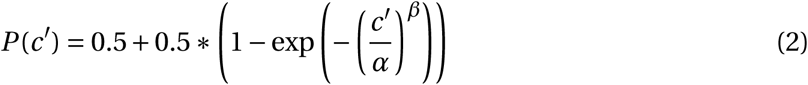

where *α* is the discrimination threshold and *β* is the psychometric order. The discrimination threshold *α* defines the coherence level that yields 81.6% correct (2). The psychometric order *β* defines the slope of the psychometric function at the the discrimination threshold. This fit function has been used previously to fit monkey psychometric functions (2, 3). To generate the probability of choices for each stimulus coherence level, we simulated 1,000 trials for each circuit model. The extended DDM computes the psychometric function exactly.

#### Psychophysical Kernel Paradigm

The psychophysical kernel (PK) paradigm is based on the experimental task design of Nienborg & Cumming (4). Stimuli are presented for 2 s, randomly sampled from a uniform distribution over coherence levels of {±6.4%, ±12.8%, ±25.6%} for each 0.05-s time bin. The psychophysical matrix (*M_PK_*) is computed as the difference in probabilities between the two choices for each coherence level at each stimulus time, normalized by the magnitude of the corresponding coherence level. The psychophysical matrix is then averaged over coherence levels to form the psychophysical kernel (PK), also known as a choice-triggered average of the stimulus. A larger PK weight *w_PK_* for a given stimulus time corresponds to a larger impact of stimuli presented at that time. Mathematically, for trials with choices to options *A* and *B* (*x_A_* and *x_B_*):

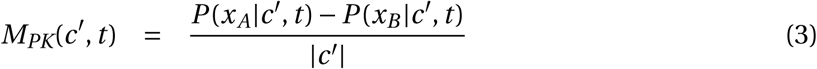

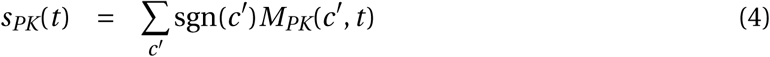

where sgn() is the sign function, returning ±1 respectively for positive and negative inputs. For this paradigm, we simulated 200,000 trials for each circuit model, and 100,000 trials for each extended DDM.

#### Pulse Paradigm

In addition to a constant 2-s stimulus of coherence levels as used in the standard task paradigm, a pulse of ±15% coherence strength and 0.1-s duration is applied at various onset times. For each pulse onset time, the psychometric function is then fitted according to

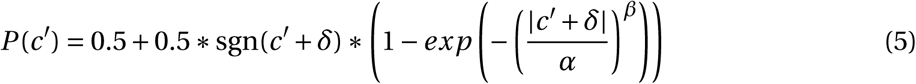

where *α*, *β*, and *δ* are the threshold, order, and shift of the psychometric function. This fit function is the same as Equation 2 but extended to include a shift *δ*, reflecting the impact of the pulse. For each circuit model, we simulated 1,750 trials, for each (15%-coherence) pulse onset time and each coherence level from the set {0%, ±3.2%, ±6.4%, ±12.8%, ±25.6%, ±51.2%}. The extended DDM computes the effect of the pulse exactly.

#### Duration Paradigm

The duration paradigm is the same as the standard task paradigm, but with stimulus presented for durations varying from 0.1 s to 2 s. For each stimulus duration, the psychometric function (Equation 2) is computed and fitted, yielding a duration-specific discrimination threshold. Each circuit model involves 1000 simulations, for each stimulus duration and coherence level. DDM computes the psychometric function exactly, such that no stochasticity is involved and no number of trials is needed.

## Extended Drift-Diffusion Model

The spiking circuit model is compared to and fitted by a drift-diffusion model (DDM). The control data is fitted by a standard DDM, where the decision variable *x* starts at 0 and migrates due to stimulus drift *μ* and noise *σ*. A choice *(A* or *B*) is selected if the decision variable reaches the corresponding bounds at *B* = ±1. To capture effects of elevated and lowered E/I ratio we extended DDM to include a self-coupling term *λ*, which affects the decision variable through an input term proportional to *x*. In the extended DDM, the dynamics of the decision variable are governed by the stochastic differential equation:

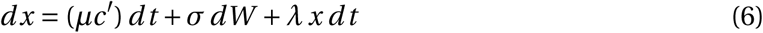

where *dW* is a Gaussian noise term of mean 0 and variance 1. *λ =* 0 corresponds to a perfect integrator used in the standard DDM. *λ* > 0, with positive self-coupling, corresponds to an unstable integrator. *λ* < 0, with negative self-coupling, corresponds to a leaky integrator.

For efficient simulation of the extended DDM, which enables fitting model parameters to spiking circuit model data and potentially experimental data, we utilized the Fokker-Planck approach. Equation (6) can be recast in terms of the probability density function (*p*(*x*, *t*)), using the Fokker-Planck equation:

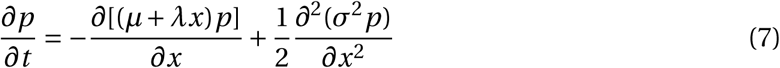

Equation (7) is solved numerically using the implicit method (staggered-mesh, absorbing boundaries) (5), with grid-size Δ*x* = 0.02 and time-step Δ*t =* 0.001. This results in a probability density function of the decision variable *x* within the boundaries *B* = ±1, as well as the probability to cross the boundaries, at each time step. After 2 s of stimulus presentation, total probability densities which crossed the upper or lower bounds are considered probabilities of either choices made, while what remains within the boundaries are deemed undecided, and are split between the two choices consistent with 2AFC.

The parameters *μ* and *σ* are determined by fitting the standard DDM (with no self-coupling strength *λ*) to the control circuit model. *λ* is then fitted separately for the other two sets of DDM parameters to elevated and lowered E/I circuits, fixing *μ* and *σ* to be the same as in the standard DDM. It was shown that fitting the models directly with all 3 parameters does not improve the fit (data not shown). From the spiking circuit data, the fitted parameter values are *μ =* 14.0, *σ =* 1.30, *λ_+_ =* 6.99, *λ_−_ =* −7.73, where *λ_+_* and *λ_−_* are the self-coupling *λ* parameter values for the elevated-E/I circuit and lowered-E/I circuit, respectively.

## Model Fitting

All fit functions are done using the method of Maximum-Likelihood Estimator (MLE). The psychometric fit functions, Equations (2) and (5), are fitted with MLE, with the maximization function for average log-likelihood:

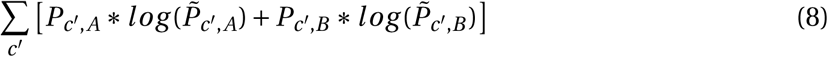

where *c′* is spanned over the coherence levels specified for each psychophysical tasks. From the circuit model or extended DDM, *P*_*c′,A*_ and *P_c′,B_* are the probabilities to select choices *A* or *B*, respectively, for each coherence level 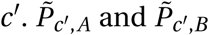 are the corresponding values for the psychometric function being fit to the model data.

The extended DDM model parameters are obtained by fitting to the statistics of internal DM-related neural trajectories of spiking circuit model, with the maximization function:

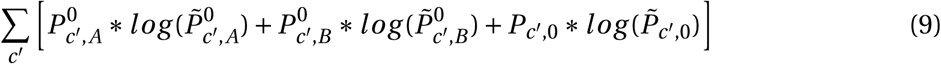

where *c′* is spanned over the coherence levels specified for the standard task paradigm. From the spiking circuit simulations, 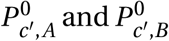 are the probabilities that population *A* or *B* crosses the firing-rate threshold, and *P*_*c′*,0_ is the indecision probability, for each coherence *c′*. From the extended DDM being fitted to the circuit data, 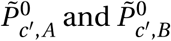 are the probabilities that the decision variable crosses the corresponding bound, and 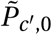 is the indecision probability, for each coherence *c′*.

## Simulation Codes

The spiking circuit model was implemented using the Python-based Brian neural simulator (6). The Fokker-Planck solver for the extended DDM was implemented in custom-written Python code. All simulation codes are available from the authors upon request.

---

**Figure S1:**
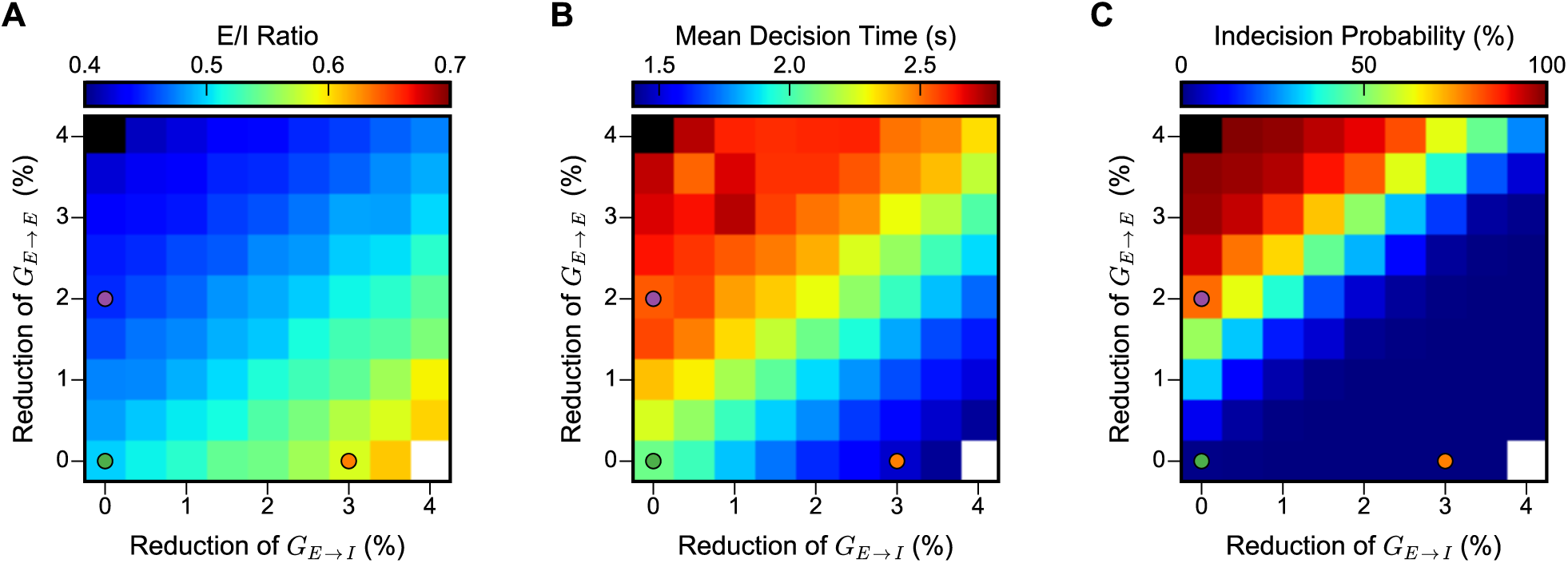
The parameter space of NMDAR hypofunction highlights the importance of E/I balance for DM function. **(A)** E/I ratio varies as a function of the perturbations of NMDAR conductance on inhibitory interneurons (*G_E→I_*) and on excitatory neurons (*G_E→E_*). E/I balance can be maintained with a proportional decrease in both *G_E→I_* and *G_E→E_*. **(B)** Mean decision time varies with E/I ratio as the critical effective parameter. **(C)** Indecision probability varies with E/I ratio as the critical effective parameter. Other conventions are as in **Figure 2** in the main text.

**Figure S2:**
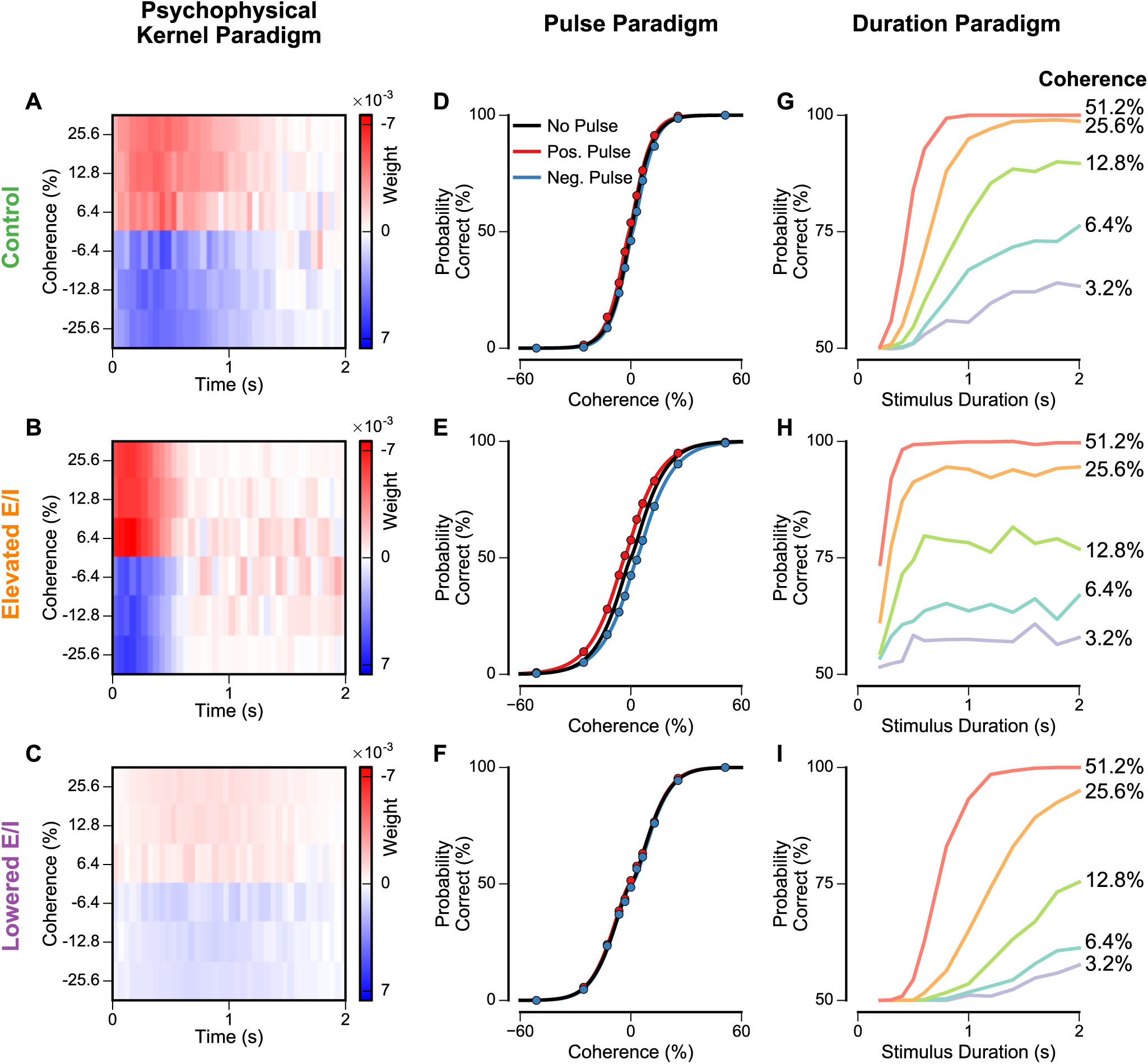
Extended circuit model behaviors for the psychophysical kernel, pulse, and duration paradigms. **(A–C)** In the psychophysical kernel paradigm, psychophysical matrices for the control, elevated-E/I, and lowered-E/I circuits. The value of each matrix element is the difference in probabilities for the two choices, for the corresponding time and coherence level, normalized by the magnitude of the coherence level. The matrix is then averaged over coherence levels to produce the psychophysical kernel (**Figure 3A** in the main text). **(D–F)** In the pulse paradigm, psychometric functions, with no pulse (black), positive pulse (red), and negative pulse (blue), for the three E/I circuits. The pulse onset time in this example is 0 s. Note that the unstable integrator has the largest shift. Simulated data (points) are fitted with Equation 5 (shown as lines). **(G–I)** In the duration paradigm, chronometric functions, the probability of a choice for option *A* as a function of stimulus duration, at various coherence levels, for the three E/I circuits.

**Figure S3:**
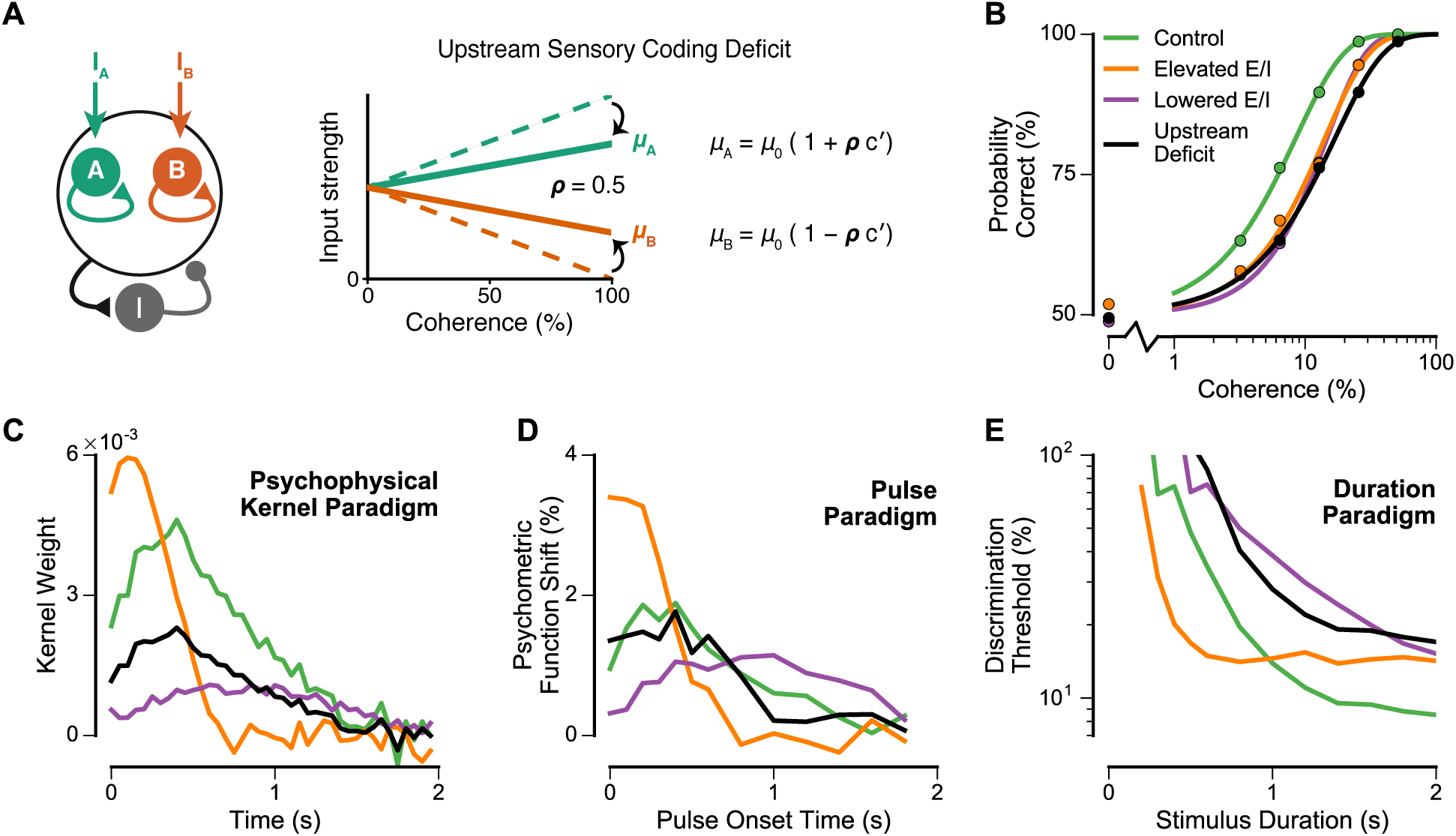
Effects of upstream deficit in sensory coding on DM performance in all four task paradigms. **(A)** Implementation of upstream sensory coding deficit in the control circuit (black lines in **Figure 3B-E**). The magnitude of this deficit is set by the parameter *ρ* which multiplicatively scales the dependence of the input firing rate to each neural population (*μ_A,B_*) on stimulus coherence (*c′*). For circuits without the upstream deficit, we use the default value *ρ =* 1. For the circuit with an upstream deficit, we set *ρ =* 0.5. **(B)** Standard task paradigm. The upstream-deficit circuit has similarly impaired performance as the elevated- and lowered- E/I circuits. **(C)** Psychophysical kernel paradigm. For the upstream-deficit circuit, the PK follows the same time course as control, but with downscaled magnitude, unlike the elevated- and lowered-E/I circuits which have altered time courses. **(D)** Pulse paradigm. For the upstream-deficit circuit, the shift curve is approximately the same as control, unlike the elevated- and lowered-E/I circuits which have different dependences on onset time. **(E)** Duration paradigm. For the upstream-deficit circuit, the threshold is higher than control but plateaus at approximately the same duration, unlike the elevated- and lowered-E/I circuits which plateau at different durations.

**Figure S4:**
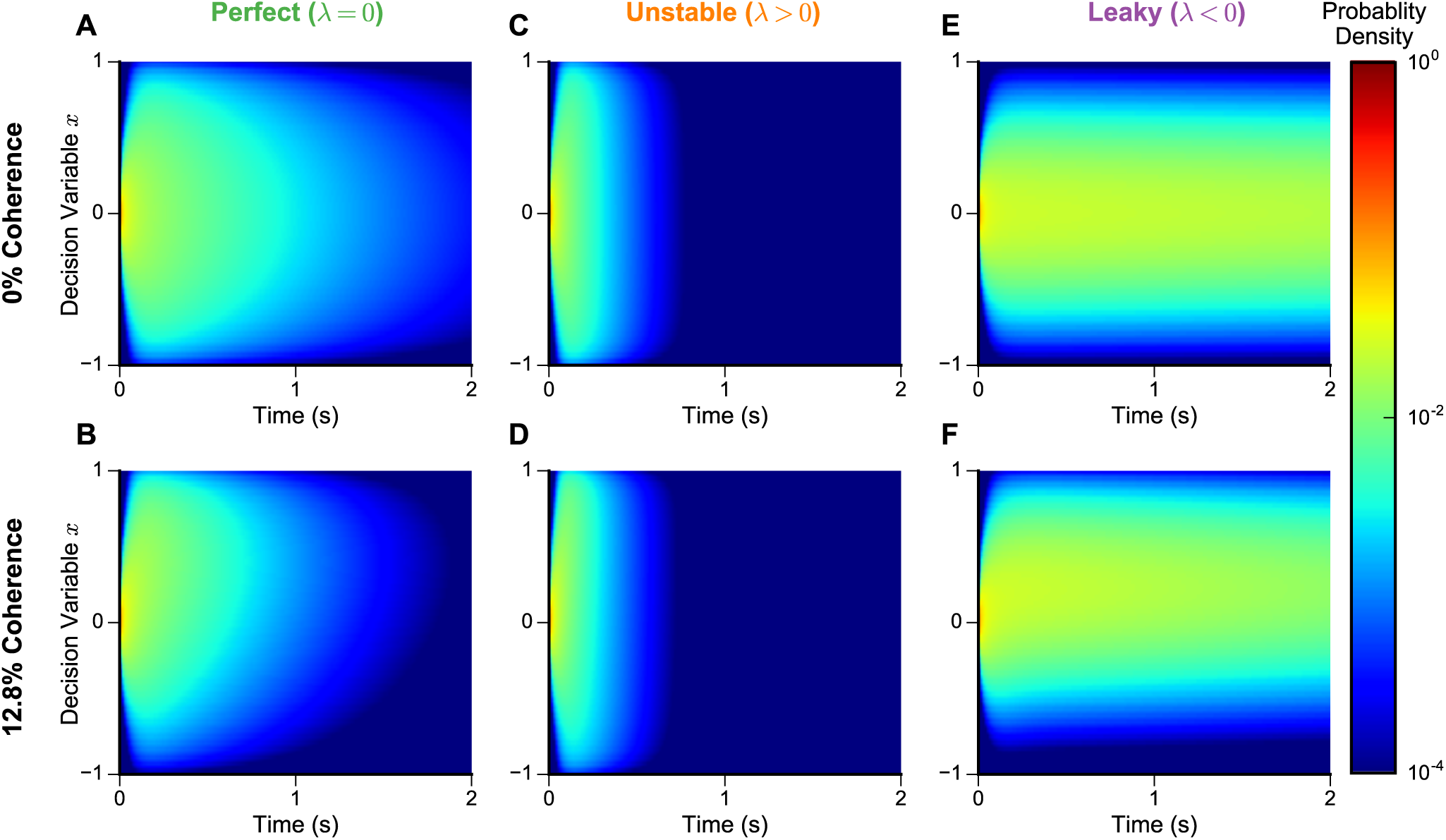
Probability density function, over time and decision variable, in the extended DDM, respectively with 0% (top) and 12.8% (bottom) coherences, for perfect *(λ =* 0) (**A**, **B**), unstable (**C**, **D**), and leaky (**E**, **F**) integrators. Parameter values used are those determined by fitting the spiking circuit model. For the unstable integrator, fitted to the elevated-E/I circuit, *λ =* 7.0 s^-1^, corresponding to a time constant of unstable growth of *τ =* 0.14 s. For the leaky integrator, fitted to the lowered-E/I circuit, *λ =* −7.7 s^−1^, corresponding to a time constant of leak of *τ =* 0.13 s.

**Figure S5:**
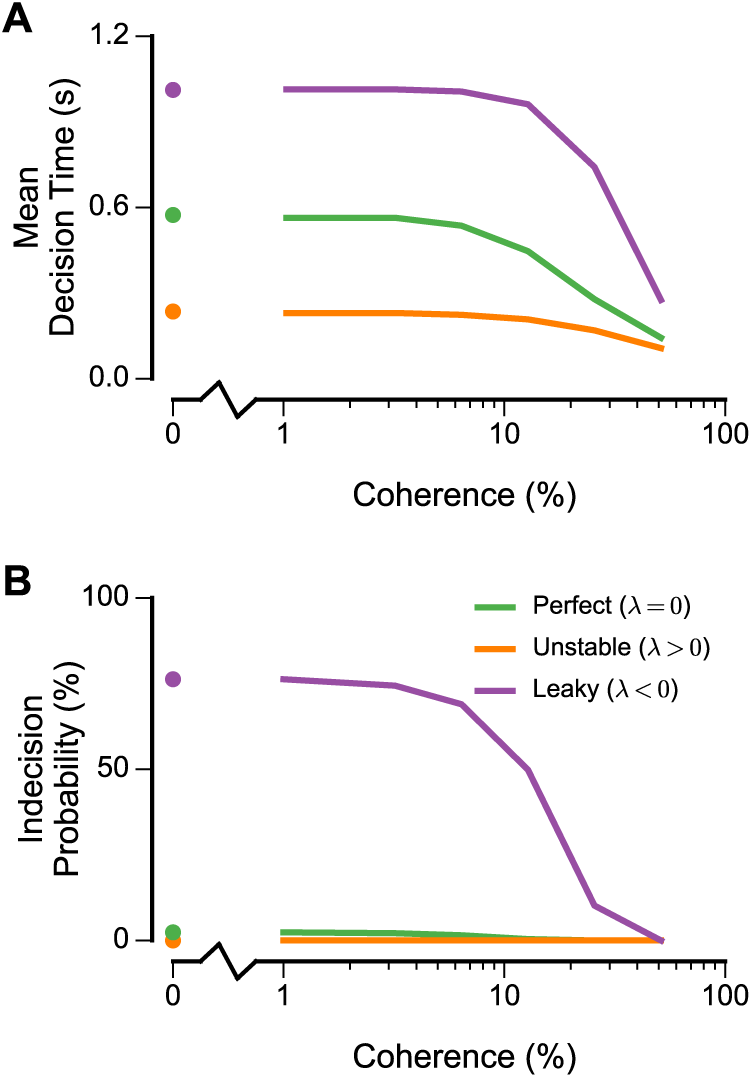
**(A)** Mean decision time of the three extended DDMs (analogue of **Figure 1D** for spiking circuit models in the main text). **(B)** Indecision probabilities of the three extended DDM (analogue of **Figure 1E** for spiking circuit models in the main text).

**Figure S6:**
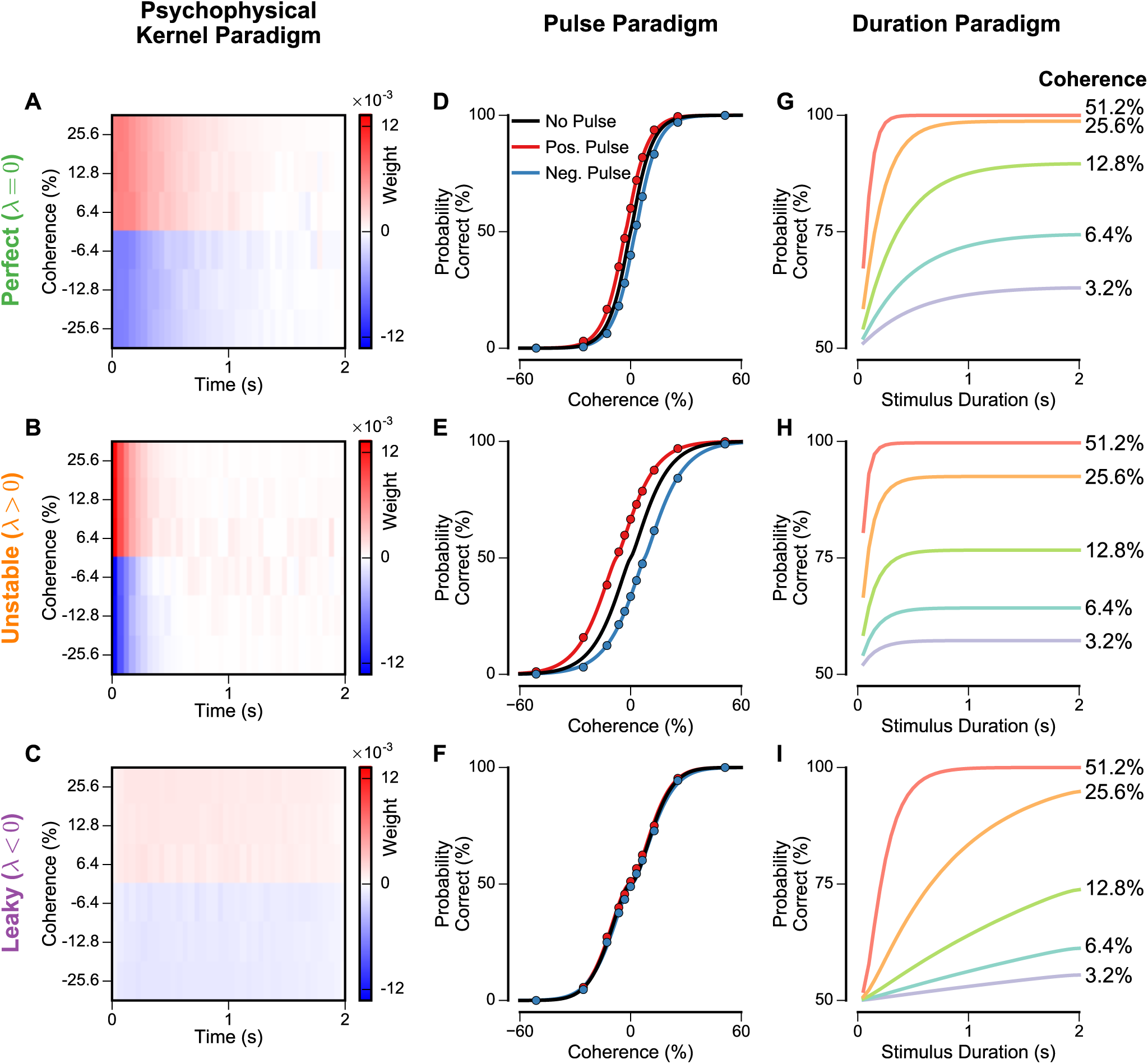
Extended DDM behaviors for the psychophysical kernel, pulse, and duration paradigms (analogue of **Figure S2** for spiking circuit models). Parameter values used are those determined by fitting the spiking circuit model. **(A–C)** In the psychophysical kernel paradigm, psychophysical matrices for the perfect, unstable, and leaky DDM. The matrix is then averaged over coherence levels to produce the psychophysical kernel (**Figure 4D** in the main text). **(D–F)** In the pulse paradigm, psychometric functions, with no pulse (black), positive pulse (red), and negative pulse (blue). The pulse onset time in this example is 0 s. Note that the unstable integrator has the largest shift. Simulated data (points) are fitted with Equation 5 (shown as lines). **(G–I)** In the duration paradigm, chronometric functions, the probability for a choice for option *A* as a function of stimulus duration, at various coherence levels.

**Figure S7:**
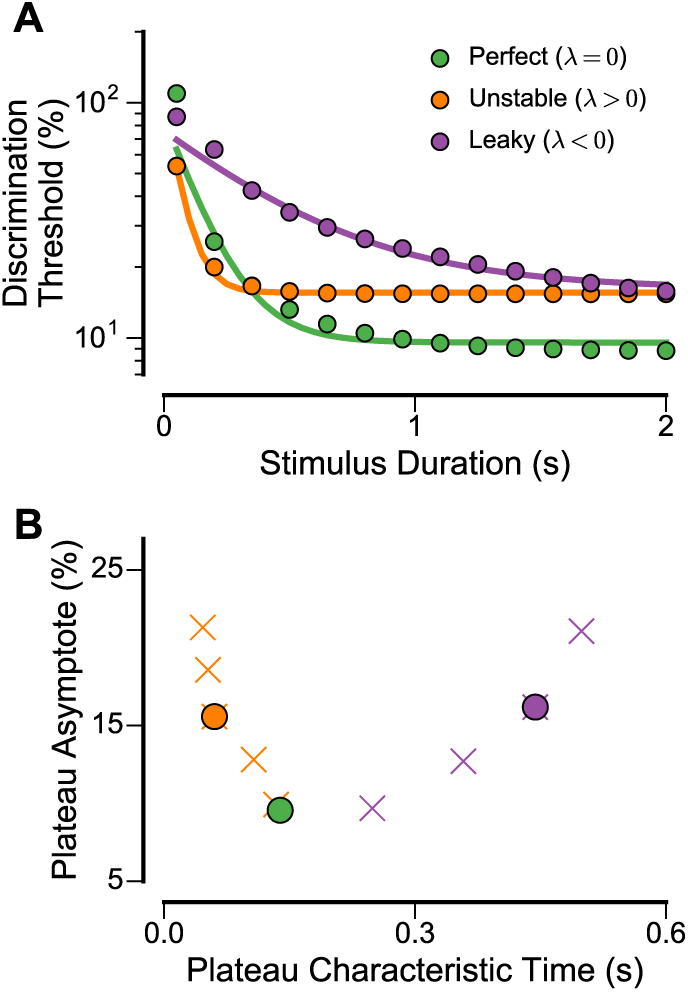
**(A)** The discrimination threshold as a function of stimulus duration for the three extended DDM models (dots) are fitted to an exponential decay function with asymptote (lines). Note that although the exponential fit cannot capture the rapid decay of the leaky integrator at early times, it does capture the general behavior otherwise. **(B)** Fitted asymptote and time constant characterizing how discrimination threshold plateaus as a function of stimulus duration. In addition to the three extended DDM models fitted to the spiking circuit models (circles), DDM models of arbitrarily chosen self-coupling strengths *λ* are similarly computed (crosses, orange for positive and purple for negative). This demonstrates how impaired performance degrades (as shown by a higher plateau asymptote) with unstable or leaky integration.

## References

1. Anticevic A, Murray JD, Barch DM (2015): Bridging levels of understanding in schizophrenia through computational modeling. Clin Psychol Sci 3:433–459.

2. Wang XJ, Krystal JH (2014): Computational psychiatry. Neuron 84:638–54.

3. Huys QJM, Maia TV, Frank MJ (2016): Computational psychiatry as a bridge from neuroscience to clinical applications. Nat Neurosci 19:404–13.

4. Murray JD, Anticevic A, Gancsos M, Ichinose M, Corlett PR, Krystal JH, Wang XJ (2014): Linking microcircuit dysfunction to cognitive impairment: effects of disinhibition associated with schizophrenia in a cortical working memory model. Cereb Cortex 24:859–72.

5. Krystal JH, D’Souza DC, Mathalon D, Perry E, Belger A, Hoffman R (2003): NMDA receptor antagonist effects, cortical glutamatergic function, and schizophrenia: toward a paradigm shift in medication development. Psychopharmacology (Berl) 169:215–233.

6. Lisman JE, Coyle JT, Green RW, Javitt DC, Benes FM, Heckers S, Grace AA (2008): Circuit-based framework for understanding neurotransmitter and risk gene interactions in schizophrenia. Trends Neurosci 31:234–242.

7. Kehrer C, Maziashvili N, Dugladze T, Gloveli T (2008): Altered excitatory-inhibitory balance in the nmda-hypofunction model of schizophrenia. Front Mol Neurosci 1:6.

8. Nakazawa K, Zsiros V, Jiang Z, Nakao K, Kolata S, Zhang S, Belforte JE (2012): GABAergic interneuron origin of schizophrenia pathophysiology. Neuropharmacology 62:1574–1583.

9. Marin O (2012): Interneuron dysfunction in psychiatric disorders. Nat Rev Neurosci 13:107–120.

10. Gao R, Penzes P (2015): Common mechanisms of excitatory and inhibitory imbalance in schizophrenia and autism spectrum disorders. CurrMolMed 15:146–67.

11. Zikopoulos B, Barbas H (2013): Altered neural connectivity in excitatory and inhibitory cortical circuits in autism. Front Hum Neurosci 7:609.

12. Hashemi E, Ariza J, Rogers H, Noctor SC, Martínez-Cerdeño V (2016): The number of parvalbumin-expressing interneurons is decreased in the medial prefrontal cortex in autism. Cereb Cortex.

13. Yizhar O, Fenno LE, Prigge M, Schneider F, Davidson TJ, O’Shea DJ, et al. (2011): Neocortical excitation/inhibition balance in information processing and social dysfunction. Nature 477:171–8.

14. Lee E, Lee J, Kim E (2016): Excitation/inhibition imbalance in animal models of autism spectrum disorders. Biol Psychiatry.

15. Elvevåg B, Goldberg T (2000): Cognitive impairment in schizophrenia is the core of the disorder. Crit Rev Neurobiol 14:1–21.

16. Barch DM, Ceaser A (2012): Cognition in schizophrenia: core psychological and neural mechanisms. Trends Cogn Sci 16:27–34.

17. Hill EL (2004): Executive dysfunction in autism. Trends Cogn Sci 8:26–32.

18. Lai CLE, Lau Z, Lui SSY, Lok E, Tam V, Chan Q, et al. (2016): Meta-analysis of neuropsychological measures of executive functioning in children and adolescents with high-functioning autism spectrum disorder. Autism Res.

19. Shadlen MN, Newsome WT (2001): Neural basis of a perceptual decision in the parietal cortex (area lip) of the rhesus monkey. J Neurophysiol 86:1916–36.

20. Roitman JD, Shadlen MN (2002): Response of neurons in the lateral intraparietal area during a combined visual discrimination reaction time task. J Neurosci 22:9475–89.

21. Gold JI, Shadlen MN (2007): The neural basis of decision making. Annu Rev Neurosci 30:535–74.

22. Kayser AS, Buchsbaum BR, Erickson DT, D’Esposito M (2010): The functional anatomy of a perceptual decision in the human brain. J Neurophysiol 103:1179–94.

23. Kelly SP, O’Connell RG (2013): Internal and external influences on the rate of sensory evidence accumulation in the human brain. J Neurosci 33:19434–41.

24. Chen Y, Nakayama K, Levy D, Matthysse S, Holzman P (2003): Processing of global, but not local, motion direction is deficient in schizophrenia. Schizophr Res 61:215–27.

25. Chen Y, Levy DL, Sheremata S, Holzman PS (2004): Compromised late-stage motion processing in schizophrenia. Biol Psychiatry 55:834–41.

26. Chen Y, Bidwell LC, Holzman PS (2005): Visual motion integration in schizophrenia patients, their first-degree relatives, and patients with bipolar disorder. Schizophr Res 74:271–81.

27. Milne E, Swettenham J, Hansen P, Campbell R, Jeffries H, Plaisted K (2002): High motion coherence thresholds in children with autism. J Child Psychol Psychiatry 43:255–63.

28. Koldewyn K, Whitney D, Rivera SM (2010): The psychophysics of visual motion and global form processing in autism. Brain 133:599–610.

29. Butler PD, Silverstein SM, Dakin SC (2008): Visual perception and its impairment in schizophrenia. Biol Psychiatry 64:40–7.

30. Wang XJ (2002): Probabilistic decision making by slow reverberation in cortical circuits. Neuron 36:955–968.

31. Wong KF, Wang XJ (2006): A recurrent network mechanism of time integration in perceptual decisions. J Neurosci 26:1314–28.

32. Wong KF, Huk AC, Shadlen MN, Wang XJ (2007): Neural circuit dynamics underlying accumulation of time-varying evidence during perceptual decision making. Front Comput Neurosci 1:6.

33. Wang XJ (2008): Decision making in recurrent neuronal circuits. Neuron 60:215–34.

34. Krystal JH, Karper LP, Seibyl JP, Freeman GK, Delaney R, Bremner JD, et al. (1994): Sub-anesthetic effects of the noncompetitive NMDA antagonist, ketamine, in humans. Psychotomimetic, perceptual, cognitive, and neuroendocrine responses. Arch Gen Psychiatry 51:199–214.

35. Anticevic A, Gancsos M, Murray JD, Repovs G, Driesen NR, Ennis DJ, et al. (2012): Nmda receptor function in large-scale anticorrelated neural systems with implications for cognition and schizophrenia. Proc Natl Acad Sci USA 109:16720–5.

36. Kiani R, Hanks TD, Shadlen MN (2008): Bounded integration in parietal cortex underlies decisions even when viewing duration is dictated by the environment. J Neurosci 28:3017–29.

37. Nienborg H, Cumming BG (2009): Decision-related activity in sensory neurons reflects more than a neuron’s causal effect. Nature 459:89–92.

38. Wimmer K, Compte A, Roxin A, Peixoto D, Renart A, de la Rocha J (2015): Sensory integration dynamics in a hierarchical network explains choice probabilities in cortical area mt. Nat Commun 6:6177.

39. Raposo D, Kaufman MT, Churchland AK (2014): A category-free neural population supports evolving demands during decision-making. Nat Neurosci 17:1784–92.

40. Huk AC, Shadlen MN (2005): Neural activity in macaque parietal cortex reflects temporal integration of visual motion signals during perceptual decision making. J Neurosci 25:10420–36.

41. Brunton BW, Botvinick MM, Brody CD (2013): Rats and humans can optimally accumulate evidence for decision-making. Science 340:95–8.

42. Smith PL, Ratcliff R (2004): Psychology and neurobiology of simple decisions. Trends Neurosci 27:161–8.

43. Bogacz R, Brown E, Moehlis J, Holmes P, Cohen JD (2006): The physics of optimal decision making: a formal analysis of models of performance in two-alternative forced-choice tasks. Psychol Rev 113:700–65.

44. Britten KH, Shadlen MN, Newsome WT, Movshon JA (1992): The analysis of visual motion: a comparison of neuronal and psychophysical performance. J Neurosci 12:4745–65.

45. Kim S, Lee D (2011): Prefrontal cortex and impulsive decision making. Biol Psychiatry 69:1140–6.

46. Evans SL, Averbeck BB, Furl N (2015): Jumping to conclusions in schizophrenia. Neuropsychiatr Dis Treat 11:1615–24.

47. Gruber AJ, Calhoon GG, Shusterman I, Schoenbaum G, Roesch MR, O’Donnell P (2010): More is less: a disinhibited prefrontal cortex impairs cognitive flexibility. J Neurosci 30:17102–10.

48. Jocham G, Hunt LT, Near J, Behrens TEJ (2012): A mechanism for value-guided choice based on the excitation-inhibition balance in prefrontal cortex. Nat Neurosci 15:960–1.

49. Ueltzhöffer K, Armbruster-Genç DJN, Fiebach CJ (2015): Stochastic dynamics underlying cognitive stability and flexibility. PLoS Comput Biol 11:e1004331.

50. Yoon JH, Grandelis A, Maddock RJ (2016): Dorsolateral prefrontal cortex GABA concentration in humans predicts working memory load processing capacity. J Neurosci 36:11788–11794.

51. Platt JR (1964): Strong inference: Certain systematic methods of scientific thinking may produce much more rapid progress than others. Science 146:347–53.

52. Carter OL, Pettigrew JD, Burr DC, Alais D, Hasler F, Vollenweider FX (2004): Psilocybin impairs high-level but not low-level motion perception. Neuroreport 15:1947–51.

53. Durstewitz D, Seamans JK (2008): The dual-state theory of prefrontal cortex dopamine function with relevance to catechol-o-methyltransferase genotypes and schizophrenia. Biol Psychiatry 64:739–49.

54. Cano-Colino M, Almeida R, Gomez-Cabrero D, Artigas F, Compte A (2014): Serotonin regulates performance nonmonotonically in a spatial working memory network. Cereb Cortex 24:2449–63.

55. Hawkins GE, Forstmann BU, Wagenmakers EJ, Ratcliff R, Brown SD (2015): Revisiting the evidence for collapsing boundaries and urgency signals in perceptual decision-making. J Neurosci 35:2476–84.

56. Usher M, McClelland JL (2001): The time course of perceptual choice: the leaky, competing accumulator model. Psychol Rev 108:550–92.

57. Roxin A, Ledberg A (2008): Neurobiological models of two-choice decision making can be reduced to a one-dimensional nonlinear diffusion equation. PLoS Comput Biol 4:e1000046.

58. Glaze CM, Kable JW, Gold JI (2015): Normative evidence accumulation in unpredictable environments. Elife 4:e08825.

59. Cheadle S, Wyart V, Tsetsos K, Myers N, de Gardelle V, Herce Castañón S, Summerfield C (2014): Adaptive gain control during human perceptual choice. Neuron 81:1429–41.

60. Wiecki TV, Poland J, Frank MJ (2015): Model-based cognitive neuroscience approaches to computational psychiatry clustering and classification. Clin Psychol Sci 3:378–399.

61. Vinogradov S, Poole JH, Willis-Shore J, Ober BA, Shenaut GK (1998): Slower and more variable reaction times in schizophrenia: what do they signify? Schizophr Res 32:183–90.

62. Ngan ET, Liddle PF (2000): Reaction time, symptom profiles and course of illness in schizophrenia. Schizophr Res 46:195–201.

63. Micallef J, Guillermain Y, Tardieu S, Hasbroucq T, Possamaï C, Jouve E, Blin O (2002): Effects of subanesthetic doses of ketamine on sensorimotor information processing in healthy subjects. Clin Neuropharmacol 25:101–6.

64. Blackman RK, MacDonald AW3rd, Chafee MV (2013): Effects of ketamine on context-processing performance in monkeys: a new animal model of cognitive deficits in schizophrenia. Neuropsychopharmacology 38:2090–100.

## Supplemental References

1. Wang XJ (2002): Probabilistic decision making by slow reverberation in cortical circuits. Neu-ron 36:955–968.

2. Kiani R, Hanks TD, Shadlen MN (2008): Bounded integration in parietal cortex underlies decisions even when viewing duration is dictated by the environment. J Neurosci 28:3017–29.

3. Roitman JD, Shadlen MN (2002): Response of neurons in the lateral intraparietal area during a combined visual discrimination reaction time task. J Neurosci 22:9475–89.

4. Nienborg H, Cumming BG (2009): Decision-related activity in sensory neurons reflects more than a neuron’s causal effect. Nature 459:89–92.

5. Butcher JC (2008): Numerical methods for ordinary differential equations. 2nd ed. Chichester, England: Wiley.

6. Goodman D, Brette R (2008): Brian: a simulator for spiking neural networks in python. Front Neuroinform 2:5.

